# Limits on prediction in language comprehension: A multi-lab failure to replicate evidence for probabilistic pre-activation of phonology

**DOI:** 10.1101/111807

**Authors:** Mante S. Nieuwland, Stephen Politzer-Ahles, Evelien Heyselaar, Katrien Segaert, Emily Darley, Nina Kazanina, Sarah Von Grebmer Zu Wolfsthurn, Federica Bartolozzi, Vita Kogan, Aine Ito, Diane Mézière, Dale J. Barr, Guillaume Rousselet, Heather J. Ferguson, Simon Busch-Moreno, Xiao Fu, Jyrki Tuomainen, Eugenia Kulakova, E. Matthew Husband, David I. Donaldson, Zdenko Kohút, Shirley-Ann Rueschemeyer, Falk Huettig

**Affiliations:** Max Planck Institute for Psycholinguistics, Nijmegen, The Netherlands; Department of Chinese and Bilingual Studies, The Hong Kong Polytechnic University, Kowloon, Hong Kong; School of Psychology, University of Birmingham, Birmingham, United Kingdom; School of Experimental Psychology, University of Bristol, Bristol, United Kingdom; School of Philosophy, Psychology and Language Sciences, University of Edinburgh, Edinburgh, United Kingdom; Institute of Neuroscience and Psychology, University of Glasgow, Glasgow, United Kingdom; School of Psychology, University of Kent, Canterbury, United Kingdom; Division of Psychology and Language Sciences, University College London, London, United Kingdom; Institute of Cognitive Neuroscience, University College London, London, United Kingdom; Faculty of Linguistics, Philology & Phonetics; University of Oxford, Oxford, United Kingdom; Department of Psychology, University of Stirling, Stirling, United Kingdom; Department of Psychology, University of York, York, United Kingdom

**Author notes:** **Corresponding author**: Mante S. Nieuwland, Max Planck Institute for Psycholinguistics, Wundtlaan 1, 6525 XD Nijmegen, The Netherlands.

## Abstract

In the last few decades, the idea that people routinely and implicitly predict upcoming words during language comprehension has turned from a controversial hypothesis to a widely-accepted assumption. Current theories of language comprehension^1–3^ posit prediction, or context-based pre-activation, as an essential mechanism occurring at all levels of linguistic representation (semantic, morpho-syntactic and phonological/orthographic) and facilitating the integration of words into the unfolding discourse representation. The strongest evidence to date for phonological pre-activation comes from DeLong, Urbach and Kutas^4^, who monitored participants’ electrophysiological brain responses as they read sentences, presented one word at a time, with expected/unexpected indefinite article + noun combinations like, “The day was breezy so the boy went outside to fly a kite/an airplane”. The sentences varied expectations (‘cloze’ probability) for a consonant- or vowel-initial noun, as determined in a sentence-completion task using other participants. Expectedly, the amplitude of the N400 event-related potential (ERP) decreased (became less negative) with increasing cloze reflecting ease of processing^5–6^. Whereas the decreased N400 at the noun could be due to its pre-activation or because high-cloze nouns are easier to integrate, crucially, N400s at the immediately-preceding article *a* or *an* showed the same relationship with cloze, i.e., encountering an indefinite article that mismatched a highly-expected word (e.g., *an* when expecting *kite*) also elicited a larger N400. This led to the claim that participants pre-activated highly-expected nouns, including their initial phonemes, based on the preceding context, with larger N400s on mismatching articles reflecting disconfirmation of this prediction.

The Delong et al. study warranted stronger conclusions than related results available at the time. Unlike previous work, it did not rely on the precursory visual-depiction of upcoming nouns, clearly de-confounded prediction and integration effects, and tested for graded phonological pre-activation of specific word form. Correspondingly, the study has been enthusiastically received as strong evidence for probabilistic phonological pre-activation, receiving over 650 citations to date and featuring in authoritative reviews^2–3^. However, there is good cause to question the soundness of the original finding (and the appropriateness of the analysis used). Attempts to replicate the critical article-effect have failed^7^. Moreover, an earlier, alternative analysis of the same data by the authors^8^ failed to reach statistical significance, but was omitted from the published report.

To obtain more definitive evidence, we conducted a direct replication study spanning ^9 laboratories (*N*_*total*_ = 334). We pre-registered one replication analysis that was faithful to the^ original, and one single-trial analysis that modeled subject- and item-level variance using linear mixed-effects models. Applying the replication analysis to our article data (Figure 1a), the original finding did not replicate: no laboratory observed a significant negative relationship between cloze and N400 at central-parietal electrodes. In contrast, the negative relationship was successfully replicated for the nouns: 6 laboratories observed such an effect and 2 laboratories observed relatively strong but non-significant effects in the expected direction (range *r* = .30 to .50). In the single-trial analysis (Fig. 1b-c), there was no statistically significant effect of cloze on article-N400s, also with stricter control for pre-article voltage levels (Supplementary Fig. 1). Crucially, there was a strong and significant cloze effect on noun-N400s (in all laboratories), which was significantly different from that on article-N400s. We observed no significant differences between laboratories for article or noun effects. Exploratory Bayesian analyses with priors based on DeLong et al. further support our conclusions (Fig. 1d, Supplementary Fig. 2). Finally, a control experiment confirmed our participants’ sensitivity to the a/an rule during online language comprehension (Supplementary Fig. 3).

**Figure 1.**
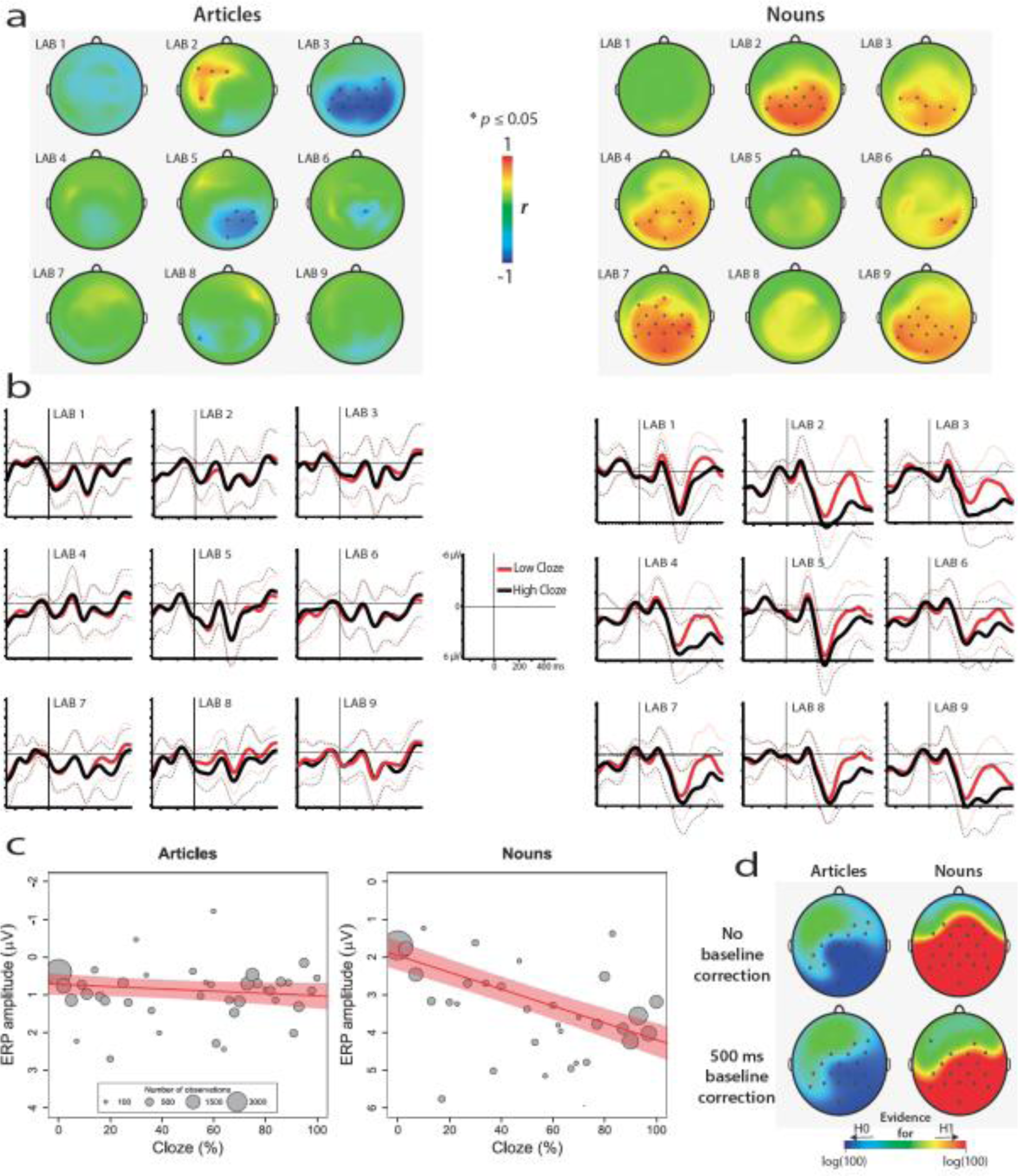
A multi-lab failure to replicate evidence for probabilistic pre-activation of phonology. (a) Pre-registered replication analysis: Pearson’s *r* correlations between ERP amplitude and article/noun cloze probability per EEG channel (* *P* < 0.05) and per laboratory. (b, c) Pre-registered single-trial analysis: (b) Grand-average ERPs elicited by relatively expected and unexpected words (cloze higher/lower than 50%) at electrode Cz, with standard deviation are shown in dotted lines, and (c) the relationship between cloze and N400 amplitude as illustrated by the mean ERP values per cloze value (number of observations reflected in circle size), along with the regression line and 95% confidence interval. A change in article cloze from 0 to 100 is associated with a change in amplitude of 0.296 µV (95% confidence interval: −.08 to .67), χ^2^(1) = 2.31, *p* = .13. A change in noun-cloze from 0 to 100 is associated with a change in amplitude of 2.22 µV (95% confidence interval: 1.75 to 2.69), χ^2^(1) = 56.5, *p* < .001. The effect of cloze on noun-N400s was statistically different from its effect on article-N400s, χ^2^(1) = 31.38, *p* < .001. (d) Bayes factor analysis associated with the replication analysis, quantifying the obtained evidence for the null hypothesis (H_0_) that N400 is not impacted by cloze, or for the alternative hypothesis (H_1_) that N400 is impacted by cloze with the size and direction of effect reported by DeLong et al. Scalp maps show the common logarithm of the replication Bayes factor for each electrode, capped at log(100) for presentation purposes. Electrodes that yielded at least moderate evidence for or against the null hypothesis (Bayes factor of ≥ 3) are marked by an asterisk. At posterior electrodes where DeLong et al. found their effects, our article data yielded strong to extremely strong evidence for the null hypothesis, whereas our noun data yielded extremely strong evidence for the alternative hypothesis (upper graphs). These results were also found when applying a 500 ms pre-word baseline correction (lower graphs).

Despite a sample size 10 times larger than the original and improved statistical analysis, we observed no statistically significant effect of cloze on article-N400s, while replicating the strong and statistically significant effect of cloze on noun-N400s^4,6^. The effect of cloze on article-N400s, if existent, must be very small to evade detection given our expansive approach. Whether such an effect would constitute convincing evidence for routine phonological pre-activation as assumed in theories of language comprehension^3^ can be questioned, but, more generally, such an effect cannot be meaningfully studied in typical small-scale studies. Consequently, current theoretical positions may be based on potentially unreliable findings and require revision. In particular, the strong prediction view that claims that pre-activation routinely occurs across all – including phonological – levels^3^, can no longer be viewed as having strong empirical support.

Our results do not constitute evidence against prediction in general. We note a lack of convincing evidence specifically for phonological pre-activation, which would have to be measured before a noun appears and unobscured by processes instigated by the noun itself.

However, our results neither support nor necessarily exclude phonological pre-activation. Unlike gender-marked articles^9^ (e.g., in Dutch or Spanish) that agree with nouns irrespective of intervening words, English a/an articles index the subsequent word, which is not always a noun. Maybe our participants did not use mismatching articles to disconfirm predicted nouns, possibly because it was not a viable strategy (American and British English corpus data show a mere 33% chance that a noun follows such articles). Perhaps a revision of the predicted meaning is required to trigger differential ERPs.

DeLong et al. recently described filler-sentences in their experiment^10, cf. 7^, which were omitted from their original report, and were neither provided nor mentioned to us upon our request for their stimuli. DeLong used the existence of these filler-sentences to dismiss an alternative explanation of their results, namely that an unusual experimental context wherein every sentence contains an article-noun combination leads participants to strategically predict upcoming nouns. Importantly, we failed to replicate their article-effects *despite* an experimental context that could inadvertently encourage strategic prediction. Therefore, the difference between their experiment and ours cannot explain the different results, and may even strengthen our conclusions.

In sum, our findings do not support a strong prediction view involving routine and probabilistic pre-activation of phonological word form based on preceding context.

Moreover, our results further highlight the importance of direct replication, large sample size studies, transparent reporting and of pre-registration to advance reproducibility and replicability in the neurosciences.

## ONLINE METHODS

### Experimental design and materials

Nieuwland requested all original materials from DeLong et al. with the stated purpose of direct replication (personal communication, November 4 and 19, 2015), upon which DeLong et al. made available the 80 sentences described in the original study. These sentences were then adapted from American to British spelling and underwent a few minor changes to ensure their suitability for British participants. The complete set of materials and the list of changes to the original materials are available online (Supplementary Table 1 and 2). The materials were 80 sentence contexts with two possible continuations each: a more or less expected indefinite article + noun combination. The noun was followed by at least one subsequent word. All article + noun continuations were grammatically correct. Within each participant, each article + noun combination served once as the more expected continuation and the other time as the less expected continuation, in different contexts. We divided the 160 items in two lists of 80 sentences such that each list contained each noun only once. Each participant was presented with only one list (thus, each context was seen only once). One in four sentences was followed by a yes/no comprehension question, which yielded a mean comprehension accuracy of 86%. This percentage cannot be directly compared to that of DeLong et al., because new comprehension questions had to be created in the absence of the original ones (but see exploratory single-trial analysis section, for relevant information).

Article cloze and noun cloze ratings were obtained from a separate group of native speakers of English who were students at the University of Edinburgh and did not participate in the ERP experiment. They were instructed to complete the sentence fragment with the best continuation that comes to mind^1^. We obtained article cloze ratings from 44 participants for 80 sentence contexts truncated before the critical article. Noun cloze ratings were obtained by first truncating the sentences after the critical articles, and presenting two different, counterbalanced lists of 80 sentences to 30 participants each, such that a given participant only saw each sentence context with the expected or the unexpected article. The obtained values closely resemble those of the original study, with the same range (0–100% for articles and nouns), slightly lower median values (for articles and nouns, 29% and 40%, compared to 31% and 46% in the original study), but slightly higher mean values (for articles and nouns, 41% and 46%, compared to 36% and 44%). Because the sentence materials we used describe common situations that can be understood by any English speaker, and because students at the University of Edinburgh come from across the whole of the UK, we had no *a priori* expectation that cloze ratings would differ substantially across laboratories, and thus we did not obtain cloze norms from other sites. Consistently, nothing in our results suggests stronger cloze effects in University of Edinburgh students compared to other students, suggesting that our cloze norms are sufficiently representative for the other universities.

### Participants

Participants were students from the University of Birmingham, Bristol, Edinburgh, Glasgow, Kent, Oxford, Stirling, York, or volunteers from the participant pool of University College London or Oxford University, who received cash or course credit for taking part in the ERP experiment. Participant information and EEG recording information per laboratory is available online (Supplementary Table 3). We pre-registered a target sample size of 40 participants per laboratory, which was thought to give at least 32 participants (the sample size of DeLong et al.) per laboratory after accounting for data loss, as was later confirmed. Due to logistic constraints, not all laboratories reached an N of 40. Because in two labs corruption of data was incorrectly assumed before computing trial loss, these laboratories tested slightly more than 40 participants. All participants (*N* = 356; 222 women) were right-handed, native English speakers with normal or corrected-to-normal vision, between 18–35 years (mean, 19.8 years), free from any known language or learning disorder. Eighty-nine participants reported a left-handed parent or sibling.

### Procedure

After giving written informed consent, participants were tested in a single session. Written sentences were presented in the center of a computer display, one word at a time (200 ms duration, 500 ms stimulus onset asynchrony). Participants were instructed to read sentences for comprehension and answer yes/no comprehension questions by pressing hand-held buttons. The electroencephalogram (EEG) was recorded from at least 32 electrodes.

The replication experiment was followed by a control experiment, which served to detect sensitivity to the correct use of the a/an rule in our participants. Participants read 80 relatively short sentences (average length 8 words, range 5–11) that contained the same critical words as the replication experiment, preceded by a correct or incorrect article. As in the replication experiment, each critical word was presented only once, and was followed by at least one more word. All words were presented at the same rate as the replication experiment. There were no comprehension questions in this experiment. After the control experiment, participants performed a Verbal Fluency Test and a Reading Span test; the results from these tests are not discussed here. All stimulus presentation scripts are publicly available in two different software packages (E-Prime and Presentation) on https://osf.io/eyzaq.

### Data processing

Data processing was performed in BrainVision Analyzer 2.1(Brain Products, Germany). We performed one pre-registered replication analysis that followed the DeLong et al. analysis as closely as possible and one pre-registered single-trial analysis (Open Science Framework, <u>https://osf.io/eyzaq).</u> All non-pre-registered analyses are considered as exploratory. First, we interpolated bad channels from surrounding channels, and downsampled to a common set of 22 EEG channels per laboratory which were similar in scalp location to those used by DeLong et al. For one laboratory that did not have all the selected 22 channels, 12 virtual channels were created using topographic interpolation by spherical splines. We then applied a 0.01–100 Hz digital band-pass filter (including 50 Hz Notch filter), re-referenced all channels to the average of the left and right mastoid channels (in a few participants with a noisy mastoid channel, only one mastoid channel was used), and segmented the continuous data into epochs from 500 ms before to 1000 ms after word onset. We then performed visual inspection of all data segments and rejected data with amplifier blocking, movement artifacts, or excessive muscle activity. Subsequently, we performed independent component analysis^2^, based on a 1-Hz high-pass filtered version of the data, to correct for blinks, eye movements or steady muscle artefacts. After this, we automatically rejected segments containing a voltage difference of over 120 µV in a time window of 150 ms or containing a voltage step of over 50 µV/ms. Participants with fewer than 60 article trials or 60 noun trials were removed from the analysis, leaving a total of 334 participants (range across laboratories 32–42, and therefore each lab had a sample size at least as large as DeLong et al.). On average, participants had 77 article trials and 77 noun trials.

### Replication analysis

We applied a 4^th^-order Butterworth band-pass filter at 0.2–15 Hz to the segmented data, averaged trials per participant within 10% cloze bins (0–10, 11–20, etc. until 91–100), and then averaged the participant-wise averages separately for each laboratory. Because the bins did not contain equal numbers of trials (the intermediate bins contained fewest trials), like in DeLong et al., not all participants contributed a value for each bin to the grand average per laboratory. For nouns and articles separately, and for each EEG channel, we computed the correlation between ERP amplitude in the 200–500 ms time window per bin with the average cloze probability per bin.

We point out that this correlation analysis reduces an initially large pool of at least 2560 potential data points per lab (32 or more subjects who each read 80 sentences), to 10 grand-average values, by averaging N400 responses over trials within 10 cloze probability decile-bins (cloze 0–10, 11–20, et cetera), per participant and then averaging over participants, even though these bins held greatly different numbers of observations. Correlating these 10 values with the average cloze value per bin yields correlation coefficients with large confidence intervals (for example, the Cz electrode in DeLong et al. showed a statistically significant *r*-value of 0.68 with a 95% confidence interval ranging from 0.09 to 0.92). By discretizing cloze probability into deciles and not distinguishing various sources of subject-, item-, bin-, and trial-level variation, this analysis potentially compromises power.

Furthermore, treating subjects as fixed rather than random potentially inflates false positive rates, due to the confounding of the overall cloze effect with by-subject variation in the effect^3–4^. Therefore, our study also seeks to improve upon DeLong et al.’s original data analysis with a pre-registered single-trial analysis.

### Pre-registered single-trial analysis

In this analysis we did not apply the 0.2–15 Hz band-pass filter, which carries the risk of inducing data distortions^5–6^. For each trial, we performed baseline correction by subtracting the mean voltage of the −100 to 0 ms time window from the data. This common procedure corrects for spurious voltage differences before word onset, generating confidence that observed effects are elicited by the word rather than differences in brain activity that already existed before the word. Baseline correction is a standard procedure in ERP research^5^, and although it was not used or not reported in DeLong et al, it has been used in many other publications from the same lab. On the basis of a review of the published work from this lab (i.e. the Kutas Cognitive Electrophysiology Lab, we have identified the 100 pre-stimulus baseline as the most frequently used one in similar studies.

Instead of averaging N400 data for subsequent statistical analysis, we performed linear mixed effects model analysis^7^ of the single-trial N400 data, using the “lme4” package^8^ in the R software^9^. This approach simultaneously models variance associated with each subject and with each item. Using a spatiotemporal region-of-interest approach based on the DeLong et al. results, our dependent measure (N400 amplitude) was the average voltage across 6 centro-parietal channels (Cz/C3/C4/Pz/P3/P4) in the 200–500 ms window for each trial. Analysis scripts and data to run these scripts are publicly available on https://osf.io/eyzaq.

For articles and nouns separately, we used a maximal random effects structure as justified by the design^4^, which did not include random effects for ‘laboratory’ as there were only 9 laboratories, and laboratory was not a predictor of theoretical interest. *Z*-scored cloze was entered in the model as a continuous variable that had two possible values for each item (corresponding to relatively expected and unexpected words), and laboratory was entered as a deviation-coded categorical variable. We tested the effects of ‘laboratory’ and ‘cloze’ through model comparison with a χ^2^ log-likelihood test. We tested whether the inclusion of a given fixed effect led to a significantly better model fit. The first model comparison examined laboratory effects, namely whether the cloze effect varied across laboratories (cloze-by-laboratory interaction) or whether the N400 magnitudes varied over laboratory (laboratory main effect). If laboratory effects were nonsignificant, we dropped them from the analysis to simplify interpretation. For the articles and nouns separately, we compared the subsequent models below. Each model included the random effects associated with the fixed effect ‘cloze’^4^. All output *β* estimates and 95% confidence intervals (CI) were transformed from z-scores back to raw scores, and then back to the 0–100% cloze range, so that the voltage estimates represent the change in voltage associated with a change in cloze probability from 0 to 100.

Model 1: N400 ~ cloze * laboratory + (cloze | subject) + (cloze | item)

Model 2: N400 ~ cloze + laboratory + (cloze | subject) + (cloze | item)

Model 3: N400 ~ cloze + (cloze | subject) + (cloze | item)

Model 4: N400 ~ (cloze | subject) + (cloze | item)

We also tested the differential effect of cloze on article ERPs and on noun ERPs by comparing models with and without an interaction between cloze and the deviation-coded factor ‘wordtype’ (article/noun). Random correlations were removed for the models to converge.

Model 1: N400 ~ cloze * wordtype + (cloze * wordtype || subject) + (cloze * wordtype || item)

Model 2: N400 ~ cloze + wordtype + (cloze * wordtype || subject) + (cloze * wordtype || item)

### Exploratory single-trial analyses

We noticed small ERP effects of cloze in the time window before article onset in laboratories 1, 3, 4, 6, 8 and 9, and a slow drift effect of cloze immediately at article onset in laboratory 8 (Supplementary Figures showing all electrodes are available on https://osf.io/eyzaq). We therefore performed an exploratory analysis in the 500 to 100 ms time window *before* the article, using the originally (−100 to 0 ms) baselined data, using Model 3 and 4 from the article analysis. This window covers the first 400 ms of the word that preceded the article. Because analysis in this window yielded a similar pattern as in the pre-registered analysis, we then performed exploratory analyses with longer (200 ms or 500 ms) pre-article baselines to better account for pre-article voltage levels (these windows are also often used in the Kutas laboratory). We also performed an exploratory analysis with the original baseline but an additional 0.1 Hz high-pass filter applied before baseline correction. We used this filter because it is frequently used in the Kutas laboratory and removes slow signal drift without impacting N400 activity (which has a higher-frequency spectrum)^5–6^. The results of these exploratory analyses did not change our conclusions and are shown in Supplementary Figure 1.

We note that our conclusions based on the single-trial analysis of the article data and noun data hold even when analyzing only those participants with an accuracy score at least as high as the lowest subject-accuracy reported in the original study (88%, reducing our sample to 161 participants, still 5 times larger than that of DeLong et al.). Although these analyses are not reported in the main text, they can be reproduced from our online data set, which includes the accuracy and stimulus list-version of each participant.

### Exploratory Bayesian analyses

Supplementing the Replication analysis, we performed a Bayes factor analysis for correlations^10^ using as prior the size and direction of the effect reported in the original study. This test was performed for each electrode separately, after collapsing the data points from the different laboratories. Because we had no articles in the 40–50 % cloze bin, there was a total of 9 and 10 data points per laboratory for the articles and nouns, respectively. Our analysis used priors estimated from the DeLong et al results matched as closely as possible to our electrode locations. A Bayes factor between 3 and 10 is considered moderate evidence, between 10–30 is considered strong evidence, 30–100 is very strong evidence, and values over 100 are considered extremely strong evidence. In addition to using a 100 ms pre-stimulus baseline, we also computed the replication Bayes factors using a 500 ms pre-stimulus time window for baseline correction. Results are shown in Figure 1.

Supplementing the single-trial analyses, we performed Bayesian mixed-effects model analysis using the brms package for R^11^, which fits Bayesian multilevel models using the Stan programming language^12^. We used a prior based on the Delong et al. observed effect size at Cz for a difference between 0% cloze and 100% cloze (1.25 㯀V and 3.75 㯀V for articles and nouns, respectively) and a prior of zero for the intercept. Both priors had a normal distribution and a standard deviation of 0.5 (given the a priori expectation that average ERP voltages in this window generally fluctuate on the order of a few microvolts; note that these units are expressed in terms of the *z*-scored cloze values, rather than the original cloze values, such that μ for the cloze prior was 0.45, which corresponds to a raw cloze effect of 1.25). We computed estimates and 95% credible intervals for each of the mixed-effects models we tested, and transformed these back into raw cloze units. The credible interval is the range of values such that one can be 95% certain that it contains the true effect, given the data, priors and the model. The results from these analyses did not change our conclusions and are shown in Supplementary Figure 2.

### Control experiment

Analysis of the control experiment involved a comparison between a model with the categorical factor ‘grammaticality’ (grammatical/ungrammatical) and a model without. Our dependent measure (P600 amplitude^13^) was the average voltage across 6 centro-parietal channels (Cz/C3/C4/Pz/P3/P4) in the 500–800 ms window for each trial. Results are shown in Supplementary Figure 3.

Model 1: P600 ~ grammaticality + (grammaticality | subject) + (grammaticality | item)

Model 2: P600 ~ (grammaticality | subject) + (grammaticality | item)

## Author contributions

M.S.N. and F.H. designed the research, M.S.N., D.J.B., G.R., and S.P.-A. planned the analysis. E.H., E.D., S.V.G.Z.W., F.B., V.K., A.I., S.B.-M., Z.F., E.K., S.P-A., and Z.K. collected data. M.S.N., K.S., N.K., G.R., H.J.F., J.T., E.M.H., D.I.D., and S.R supervised data collection. M.S.N. and S.P.-A. analyzed the data. M.S.N. drafted the manuscript and received comments from S.P.-A., N.K., K.S., D.J.B., H.J.F., E.M.H, and F.H.

## Acknowledgements

This work was partly funded by ERC Starting grant 636458 to H.J.F.

---

**Supplementary Figure 1.**
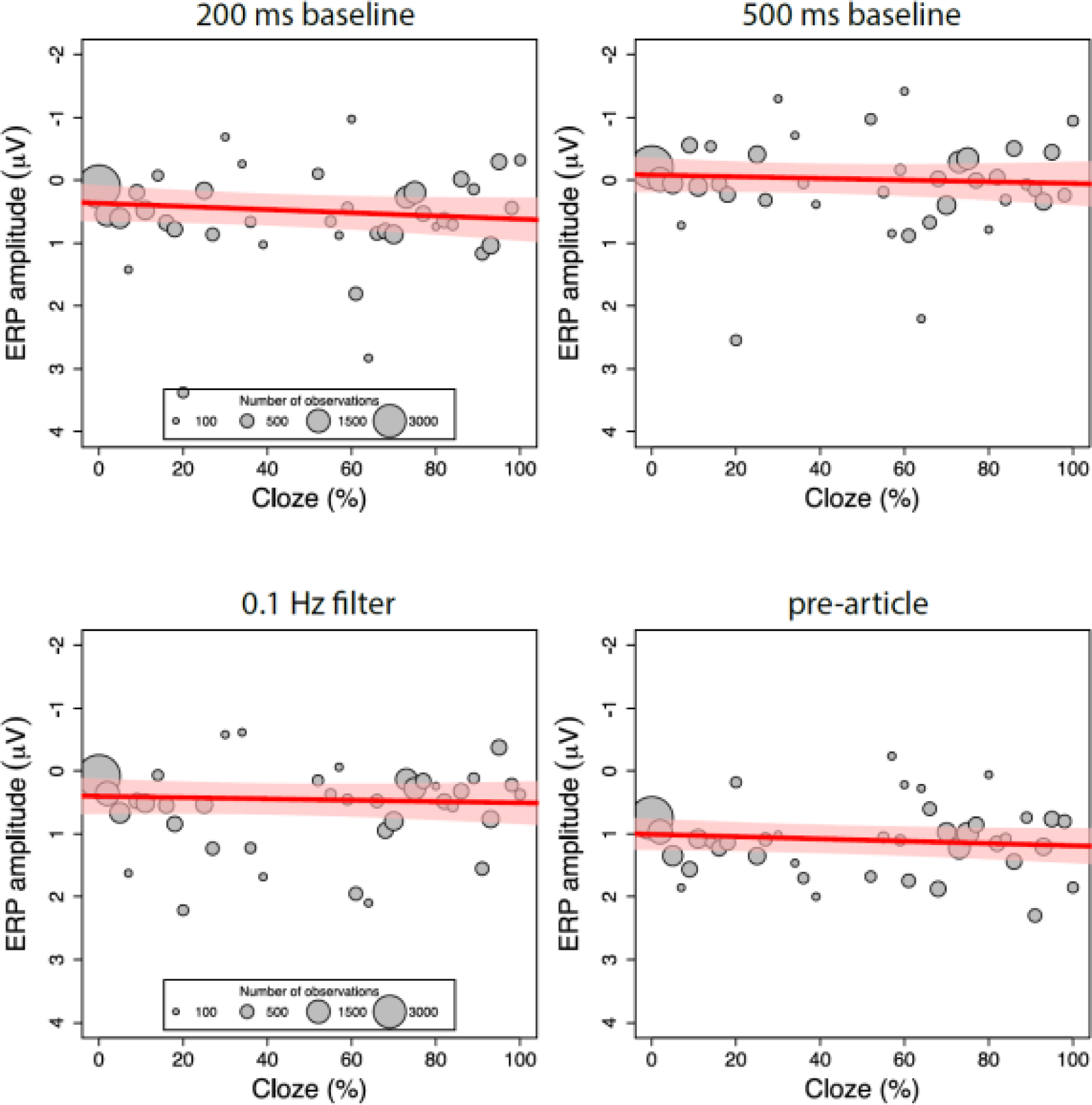
Exploratory single-trial analyses: The relationship between cloze and ERP amplitude as illustrated by the mean ERP values per cloze value (number of observations reflected in circle size), along with the regression line and 95% confidence interval, from four exploratory analyses. We performed tests which used longer baseline time windows (200 ms, upper left panel; 500 ms, upper right panel) to better control for pre-article voltage levels, or which used the pre-registered baseline and applied a 0.1 Hz high-pass filter (lower left panel) to better control for slow signal drift (while presumably not affecting N400 activity). All three tests reduced the initially observed effect of article-cloze (200 ms baseline, β= .25, CI [−.12, .62], χ^2^(1) = 1.35, *p* = .19; 500 ms baseline, β= .14, CI [−.25, .53], χ^2^(1) = 0.46, *p* = .50; 0.1 Hz filter: β= 0.09, CI [−.22, .41], χ^2^(1) = 0.33, *p* = .56). An analysis in the 500 to 100 ms time window *before* article-onset (lower right panel) revealed a non-significant effect of cloze that resembled the pattern observed *after* article-onset, β= .16, CI [−.07, .39], χ^2^(1) = 1.82, *p* = .18. Combined, these results suggest that the results obtained with the pre-registered analysis at least partly reflected the effects of slow signal drift that existed before the articles were presented.

**Supplementary Figure 2.**
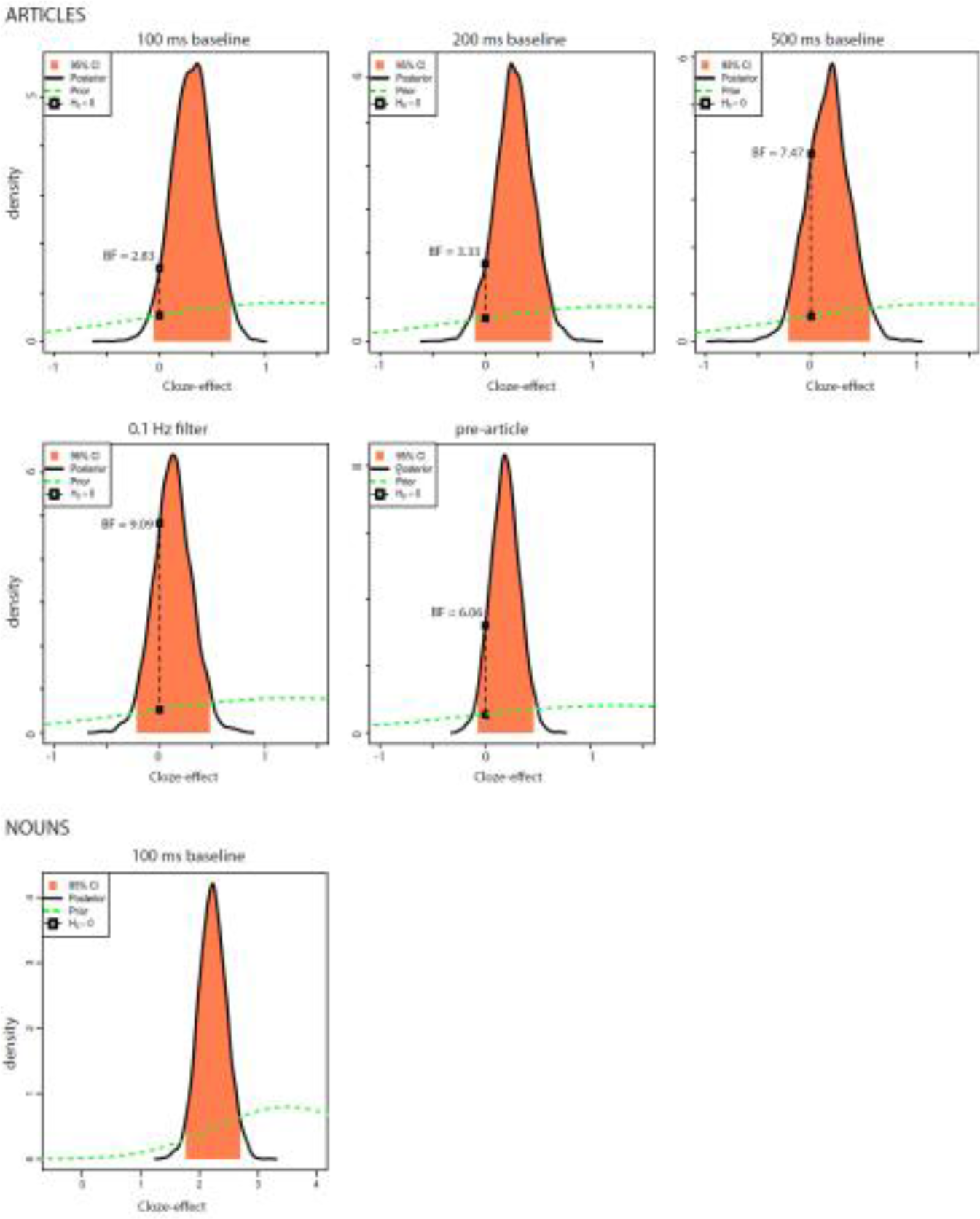
Results from exploratory Bayesian mixed-effects model analyses, represented by posterior distributions for the effect of cloze on ERP amplitudes in the N400 window. The x-axis shows cloze effect sizes (i.e., changes in microvolts associated with an increase from 0% cloze probability to 100% cloze probability). The black line indicates the posterior distribution of effects; higher values of the posterior density at a given effect size indicate higher probability that this is the true effect size in the population. The peak of the posterior distribution roughly corresponds to the point estimate of the effect size (the regression coefficient) fitted from the Bayesian mixed effect model, i.e., the most likely value of the true effect size. The middle 95% of the posterior distribution, shaded in pink, corresponds to a two-tailed 95% credible interval for the effect size—i.e., an interval that we can be 95% confident contains the true effect. The green dotted line indicates the prior distribution (i.e., our expectation about where the true effect would lie before the data were collected), which is centered on 1.25μV, the effect observed by Delong and colleagues (2005). The black connected dots illustrate the ratio between the posterior and prior distribution (i.e., the Bayes Factor) at the effect size of 0μV; for example, a Bayes Factor of 4 suggests we can be 4 times more certain that the true effect is zero after having conducted this experiment than before, or, in other words, that the data increased our confidence in the null effect of zero fourfold. We performed these analyses for each of the linear mixed-effects model analysis we performed. We note that in all the article-analyses, the posterior probability of the estimated effect being greater than zero is around 80 or 90%, but this is also the case for the pre-stimulus variable, suggesting that the observed patterns arise before the articles are seen. In none of our article-analyses did zero lie outside the obtained credible interval, whereas for the nouns, zero lay outside the credible interval. These results are consistent with a failure to replicate the DeLong et al. article-effect and successful replication of the noun-effect.

**Supplementary Figure 3.**
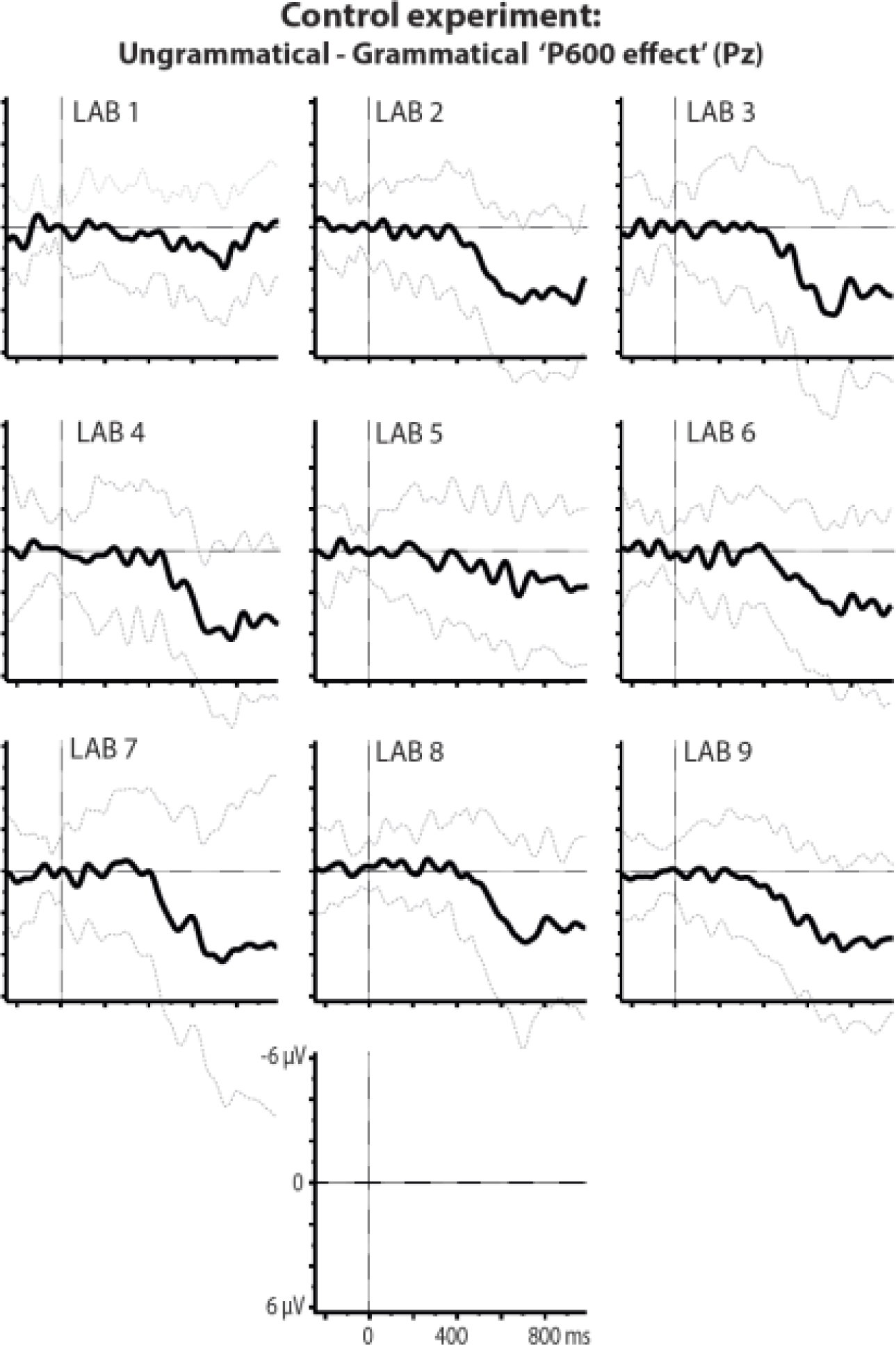
P600 effects at electrode Pz per lab associated with flouting of the English a/an rule in the control experiment. Plotted ERPs show the grand-average difference waveform and standard deviation for ERPs elicited by ungrammatical expressions (‘an kite’) minus those elicited by grammatical expressions (‘a kite’). This control experiment followed in the same experimental session as the main experiment and was carried to rule out that an observed lack of a statistically significant, article-elicited prediction effect in the main experiment reflected a general insensitivity of our participants to the a/an rule. In each laboratory, nouns following incorrect articles elicited a late positive-going waveform compared to nouns following correct articles, starting at about 500 ms after word onset and strongest at parietal electrodes. This standard P600 effect was confirmed in a single-trial analysis, χ^2^(1) = 83.09, *p* < .001, and did not significantly differ between labs, χ^2^(8) = 8.98, *p* = .35.

## Method REFERENCES

1 Taylor, W.L. Journalism Quart. 30, 415–433 (1953).

2 Jung, T.P., et al. Psychophysiology. 37, 163–178 (2000).

3 Clark, H.H. J. Verb. Learn. Verb. Behav. 12, 335–359 (1973).

4 Barr, D.J., Levy, R., Scheepers, C. & Tily, H.J. J. Mem. Lang. 68, 255–278 (2013).

5 Luck, S. J. MIT press, Cambridge MA, (2014).

6 Tanner, D., Morgan-Short K., & Luck, S. J. Psychophysiology. 52, 997–1009 (2015).

7 Baayen, R.H., Davidson, D.J. & Bates, D.M. J. Mem. Lang. 59, 390–412 (2008).

8 Bates, D., Maechler, M., Bolker, B., & Walker, S. R package version, 1 (2014).

9 R CoreTeam, R Foundation for Statistical Computing. Vienna, Austria. URL 〈http://www.R-project.org/〉. (2014)

10 Wagenmakers, E.J., Verhagen, J. & Ly, A. Behav. Res. Meth. 48, 413–426 (2016).

11 Buerkner, P.C. R package version 1.4.0 (2016).

12 Stan Development Team. R package version 2.14.1. http://mc-stan.org. (2016)

13 Osterhout, L. & Holcomb, P.J. J. Mem. Lang. 31, 785–806 (1992).

## Method References

1 Hagoort, P. Neurosci. Biobehav. Rev. doi:10.1016/j.neubiorev.2017.01.048.

2 Lau, E.F., Phillips, C. & Poeppel, D. Nature Rev. Neurosci. 9, 920–933 (2008).

3 Pickering, M.J. & Garrod, S. Behav. Brain Sci. 36, 329–347 (2013).

4 DeLong, K.A., Urbach, T.P. & Kutas, M. Nature Neurosci. 8, 1117–1121 (2005).

5 Kutas, M. & Hillyard, S.A. Science 207, 203–205 (1980).

6 Kutas, M. & Hillyard, S.A. Nature 307, 161–163 (1984).

7 Ito, A., Martin, A.E., & Nieuwland, M.S. Lang. Cogn. Neurosci. (2017).

8 DeLong, K.A. (2009). Doctoral dissertation. San Diego: University of California.

9 Van Berkum, J.J., Brown, C.M., Zwitserlood, P., Kooijman, V. & Hagoort, P. J. Exp. Psychol. Learn. Mem. Cog. 31, 443–467 (2005).

10 DeLong, K.A., Urbach, T.P. & Kutas, M. Lang. Cogn. Neurosci. (2017).

